# Condensation of Rubisco into a proto-pyrenoid in higher plant chloroplasts

**DOI:** 10.1101/2020.10.26.354332

**Authors:** Nicky Atkinson, Yuwei Mao, Kher Xing Chan, Alistair J. McCormick

**Affiliations:** SynthSys & Institute of Molecular Plant Sciences, School of Biological Sciences, King’s Buildings, University of Edinburgh, EH9 3BF, UK; Carl R. Woese Institute for Genomic Biology, University of Illinois at Urbana-Champaign, 1206 Gregory Drive, Urbana IL 61801

## Abstract

Photosynthetic CO_2_ fixation in plants is limited by the inefficiency of the CO_2_-assimilating enzyme Rubisco (D-ribulose-1,5-bisphosphate carboxylase/ oxygenase)^1–3^. In plants possessing the C3 pathway, which includes most major staple crops, Rubisco is typically evenly distributed throughout the chloroplast stroma. However, in almost all eukaryotic algae Rubisco aggregates within a microcompartment known as the pyrenoid, in association with a CO_2_-concentrating mechanism that improves photosynthetic operating efficiency under conditions of low inorganic carbon^4^. Recent work has shown that the pyrenoid matrix is a phase-separated, liquid-like condensate^5^. In the alga *Chlamydomonas reinhardtii*, condensation is mediated by two components: Rubisco and the linker protein EPYC1 (Essential Pyrenoid Component 1)^6,7^. Here we show that expression of mature EPYC1 and a plant-algal hybrid Rubisco leads to spontaneous condensation of Rubisco into a single phase-separated compartment in Arabidopsis chloroplasts, with liquid-like properties similar to a pyrenoid matrix. The condensate displaces the thylakoid membranes and is enriched in hybrid Rubisco containing the algal Rubisco small subunit required for phase separation. Promisingly, photosynthetic CO_2_ fixation and growth is not impaired in stable transformants compared to azygous segregants. These observations represent a significant initial step towards enhancing photosynthesis in higher plants by introducing an algal CO_2_-concentrating mechanism, which is predicted to significantly increase the efficiency of photosynthetic CO_2_ uptake^8,9^.

## Main Text

Rubisco catalyses net CO_2_-fixation in all photosynthetic organisms but also reacts with oxygen, leading to a wasteful loss of previously fixed CO_2_, nitrogen and energy^1^. In plants with the C3 photosynthetic pathway (e.g. rice, wheat and soybean), Rubisco has a relatively slow turnover rate and operates under sub-optimal CO_2_ concentrations of less than half that required for maximal rates of carboxylation^10–12^. Improving the operating efficiency of Rubisco remains a key strategy to increase productivity in C3 crops^13^. This is an important goal given the steep trajectory of crop yield improvements required to meet the demands of our rising global population^14^.

Most unicellular eukaryotic photosynthetic organisms and some non-vascular land plants have evolved highly efficient CO_2_-concentrating mechanisms (CCMs) that condense Rubisco into a micro compartment within the chloroplast called the pyrenoid^4,15^. The CCM functions to enrich the pyrenoid with high concentrations of CO_2_ to enhance the performance of Rubisco carboxylation and suppress oxygenation^16^. Introduction of such CCMs into C3 plants is predicted to lead to significant increases in CO_2_-fixation efficiency and biomass yield^14,17,18^. Recent work in the model green alga *Chlamydomonas reinhardtii* (hereafter Chlamydomonas) has shown that the pyrenoid is a liquid like-condensate that undergoes liquid-liquid phase separation when the CCM is active^5^. Phase separation is facilitated by multivalent interactions between the intrinsically disordered Rubisco-binding protein EPYC1 and the small subunit of Rubisco (SSU)^7^, of which residues on the α-helices of the native SSU isoforms are critical for binding to EPYC1^19,20^. Previously we have shown that Arabidopsis can assemble a functional plant-algal hybrid Rubisco complex, with the native large subunit of Rubisco (LSU) and an SSU from Chlamydomonas, which has similar kinetic properties to wild-type (WT) Arabidopsis Rubisco^21^. However, expression of the full coding sequence of EPYC1 in Arabidopsis with plant-algal hybrid Rubisco did not result in Rubisco condensation^19^. Evidence from Chlamydomonas and *in vitro* studies suggested that an appropriate stoichiometric balance between EPYC1 and Rubisco, and possibly additional regulatory components, might be required for efficient phase separation^4,5,7,22^.

We sought to test whether high levels of expression of a mature form of EPYC1, could lead to phase separation in a higher plant chloroplast. EPYC1 was truncated according to the predicted transit peptide cleavage site between residues 26 (V) and 27 (A)^19^. A dual GFP expression system was developed to achieve high levels of EPYC1 expression and a favourable stoichiometry with Rubisco. This consisted of a binary vector containing two gene expression cassettes, each encoding mature EPYC1 with an Arabidopsis chloroplastic signal peptide and fused to a different version of GFP (turboGFP (tGFP) or enhanced GFP (eGFP)) to avoid possible recombination events (Fig. 1a, Extended Data Fig. 1). The dual GFP construct (EPYC1-dGFP) was transformed into WT plants or the Arabidopsis *1a3b* Rubisco mutant complemented with an SSU from Chlamydomonas (S2_Cr_)^21^. The resulting transgenic plants (Ep) expressed both EPY C1:: eGFP and EPYC1::tGFP, of which the latter was generally more abundant at the protein level (Fig. 1b, Extended Data Fig. 2). Previously, immunoblots against full length EPYC1 expressed in S2_Cr_ or WT plants showed additional lower molecular weight bands indicative of proteolytic degradation^19^. Here, degradation products of the mature EPYC1 were relatively reduced when expressed in S2_Cr_ compared to WT plants.

**Figure 1.**
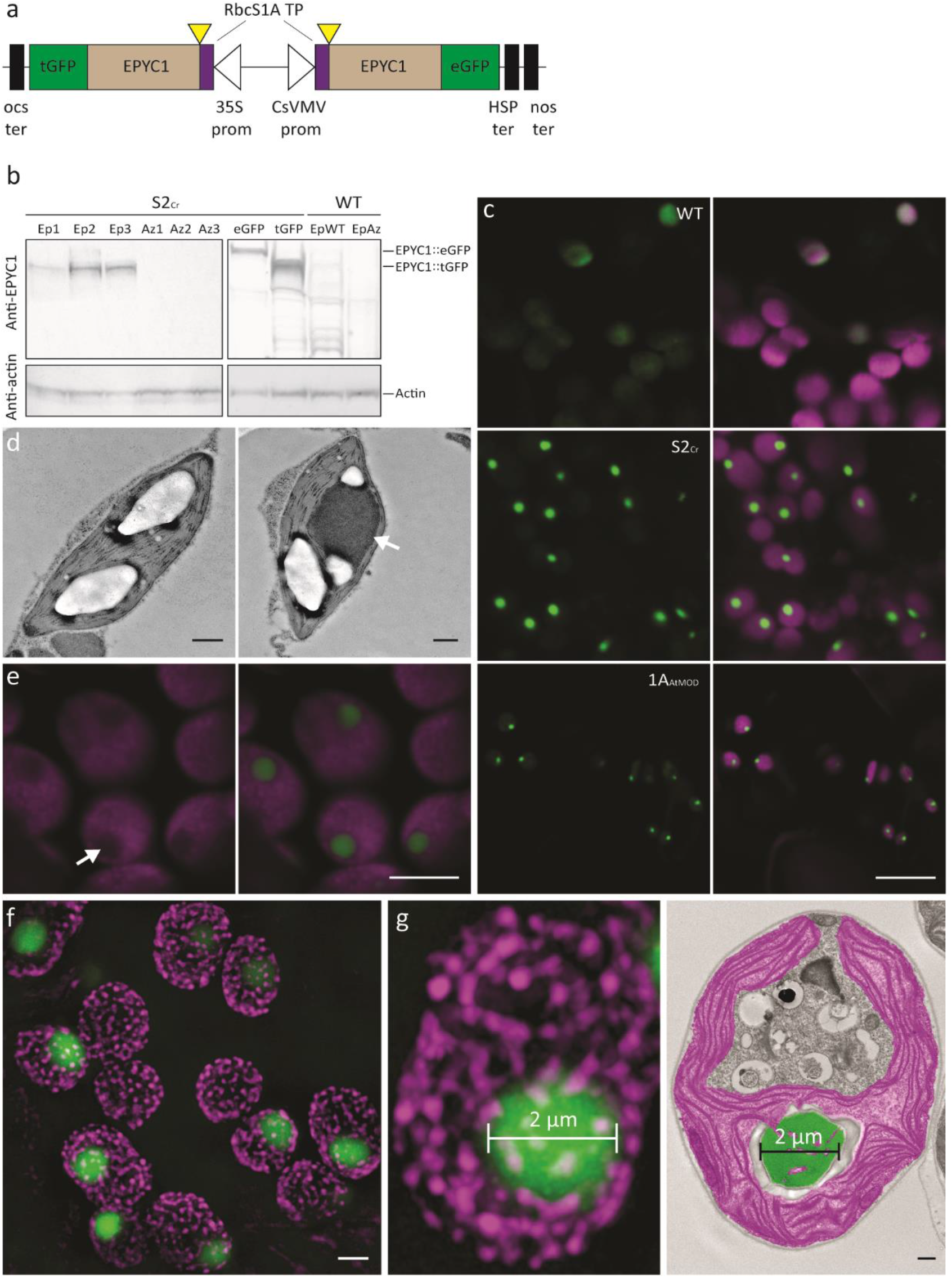
Expression of EPYC1 in the Arabidopsis line S2_Cr_ results in condensate formation. **a**, Schematic representation of the dual GFP expression system (EPYC1-dGFP) for EPYC1 truncated at amino acid residue 27 (as indicated by yellow triangles) and fused at the N-terminus to the chloroplastic transit peptide (TP) sequence of the Arabidopsis Rubisco small subunit RbcS1A. EPYC1 was further fused at the C-terminus to either enhanced GFP (eGFP) or turboGFP (tGFP), and driven by the 35S CaMV promoter (35S prom) or CsVMV promoter (CsVMV prom), respectively. For the latter expression cassette, a dual terminator system was used to increase expression^51^. **b**, EPYC1 protein levels in Arabidopsis plants as assessed by immunoblot analysis with anti-EPYC1 antibodies. Shown are three T2 S2_Cr_ transgenic plants expressing EPYC1-dGFP (Ep1-3) and azygous segregants (Az1-3), and S2_Cr_ plants transformed with only EPYC1::tGFP (55.4 kDa) or EPYC1::eGFP (63.9 kDa). Also displayed are a T2 EPYC1-dGFP WT transformant (EpWT) and azygous segregant (AzWT). Anti-actin is shown as a loading control underneath. **c**, Expression of EPYC1-dGFP in WT, S2_Cr_ and 1A_At_MOD backgrounds. Green and purple signals are GFP and chlorophyll autofluorescence, respectively. Overlapping signals are white. Scale bars = 10 μm. **d**, TEM images of chloroplasts from S2_Cr_ plants with (right) and without (left) expression of EPYC1. A white arrowhead indicates the dense dark grey area of the EPYC1 condensate. The large white structures are starch granules. Scale bars = 0.5 μm. A representative chloroplast from a wild-type plant expressing EPYC1-dGFP is shown for comparison in Extended Data Figure 3. **e**, Chlorophyll autofluorescence is reduced at the site of EPYC1-dGFP accumulation (white arrow). Scale bar = 5 μm. **f**, SIM microscopy showing EPYC1-dGFP condensates inside the chloroplast. The magenta puncta show the position of grana stacks. Light magenta puncta indicate grana stacks behind the condensate. Scale bar = 2 μm. **g**, Example comparison of condensate size (left, 2 μm) with that of a pyrenoid in Chlamydomonas (right, representative TEM image highlighting the pyrenoid in green and chloroplast in purple). Scale bar for TEM image = 0.5 μm. Images of EPYC1-dGFP condensates in the S2_Cr_ background are from line Ep3.

The fluorescence signal for EPYC1-dGFP in WT plants was distributed evenly throughout the chloroplast (Fig. 1c, Fig. 2a). In contrast, EPYC1-dGFP in the hybrid S2_Cr_ plants showed only a single dense chloroplastic signal. Transmission electron microscopy confirmed the presence of a single prominent condensed complex in the chloroplast stroma (Fig. 1d). The condensates were spherical in shape and displaced native chlorophyll autofluorescence (Fig. 1e, f), indicating that the thylakoid membrane matrix was excluded from the condensate. In protoplasts of leaf mesophyll cells, a condensate was visible in each chloroplast (Extended Data Fig. 3a), and the average size of the condensates was related to the expression level of EPYC1-dGFP (Fig. 1b, Extended Data Fig. 3b, c). We observed that the average diameter of the condensates was 1.6±0.1 μm (n=126; 42 from three individual S2_Cr_ transgenic lines), which was comparable with that of the size range measured for the Chlamydomonas pyrenoid (1.4±0.1 μm; n=55)^23^ (Fig. 1g, Extended Data Fig. 3c). The estimated volume of the condensates was 2.7±0.2 μm^3^ (*ca*. 5% of the chloroplast volume). Variations in condensate volume within individual S2_Cr_ transgenic Ep lines were not correlated with chloroplast volume, suggesting that regulation of condensate formation and size was largely independent of chloroplast morphology, and likely limited by the influence of the surrounding network stiffness (i.e. the stroma, which is densely packed with thylakoids) on the dynamics of droplet ripening^24^.

**Figure 2.**
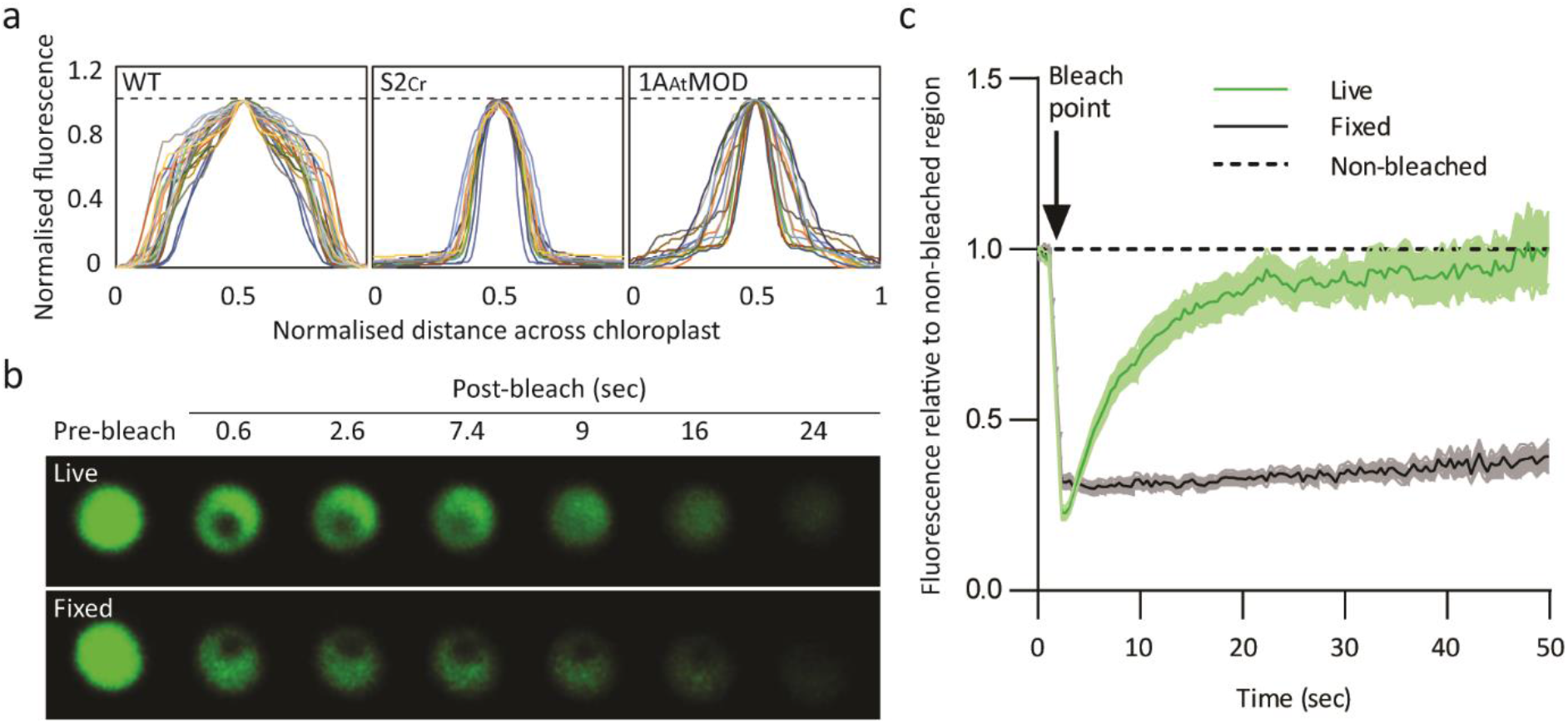
*In planta* condensates behave like liquid-liquid phase separated micro compartments. **a**, Fluorescence distribution plots of EPYC1-dGFP across the chloroplast. The intensity of the GFP fluorescence signal over the cross section of a chloroplast is shown in WT (n=28), S2_Cr_ (line Ep3, n=17) and 1A_At_MOD (n=22) backgrounds. Both GFP fluorescence and cross section values have been normalised to 1 (as indicated by the dashed line), with the highest value in the centre. **b**, Fluorescence recovery after photobleaching (FRAP) assays. Condensates are shown from live (top) and fixed (bottom) leaf tissue from S2_Cr_ transgenic line Ep3 expressing EPYC1-dGFP. **c**, Fluorescence recovery of the bleached area in relation to the non-bleached area of condensates. The mean and SEM are shown for 13-16 chloroplasts.

Condensates were also observed when EPYC1-dGFP was expressed in the Arabidopsis *1a3b* Rubisco mutant complemented with a native Arabidopsis SSU modified to contain the two α-helices necessary for pyrenoid formation from the Chlamydomonas SSU (1A_At_MOD)^20,21^ (Fig. 1c). However, condensates in the 1A_At_MOD background were less punctate (Fig. 2a), which is consistent with the lower affinity of the modified native SSU for EPYC1 observed previously in yeast two-hybrid experiments^19^. Furthermore, we confirmed that condensates were formed when either EPYC1::tGFP or EPYC1::eGFP expression cassettes were transformed into S2_Cr_ individually (Extended Data Fig. 4a). Expression of a full length (i.e. non-truncated) variant of EPYC1-dGFP in Arabidopsis chloroplasts previously did not result in phase separation^19^, which was attributed to low levels of expression and an incompatible stoichiometry between EPYC1 and Rubisco, and possible proteolytic degradation. The results here indicate that the formation of a condensate is primarily due to the expression of the mature variant of EPYC1 rather than the level of expression *per se*, and that the required stoichiometry with Rubisco is achievable *in planta*. Furthermore, the observed reduction in proteolytic degradation may be accounted for by sequestration of EPYC1 within a phase-separated compartment (Fig. 1b), which would be less accessible to large protease complexes^25^.

Next we tested if the condensates exhibited internal mixing characteristics consistent with the liquid-like behavior of pyrenoids (Fig. 2b, 2c). Fluorescence recovery after photobleaching (FRAP) assays showed that condensates in live leaf cells had similar or increased rates of rehomogenisation of EPYC1-dGFP (i.e. full recovery after 20-40 sec) compared to previous *in vitro*^7^ and *in alga*^5^ reports. The more rapid interchange in condensates compared to Chlamydomonas pyrenoids may be due to the relatively reduced availability of EPYC1 binding sites on Rubisco in the S2_Cr_ plant-algal hybrid Rubisco background^4^. In contrast, condensates in leaf tissue chemically cross-linked with formaldehyde showed no recovery after photobleaching, which is consistent with that observed for Chlamydomonas pyrenoids^5^.

To test for the presence of Rubisco, the condensates were extracted from leaf tissue and examined by immunoblot. The condensates could be pelleted by gentle centrifugation^6^, and isolated condensates were shown to be enriched in EPYC1-dGFP and Rubisco (Fig. 3a). Protein samples were probed with either a polyclonal Rubisco antibody with a greater specificity for higher plant LSU and SSUs, or an antibody raised against the RbcS2 SSU isoform of Chlamydomonas (CrRbcS2). Comparison of the resulting immunoblot data showed enrichment of the Chlamydomonas SSU in the pelleted condensate compared to native SSUs. Subsequent Coomassie staining of denatured, gel-separated extracts revealed that nearly half (49%) of Rubisco in the initial extract contained Chlamydomonas SSU, while 82% of Rubisco in the pelleted condensate contained Chlamydomonas SSU (Fig. 3b). Pelleted condensates coalesced into larger droplets when resuspended, as expected for the liquid-like behavior of EPYC1-Rubisco interactions (Fig. 3c)^7^. Immunogold analysis of chloroplast TEM images from S2_Cr_ plants expressing EPYC1-dGFP showed approximately half (54%) of all Rubisco was contained within the condensate (Fig. 3d, 3e, Extended Data Fig. 4b). Consistent with Coomasie staining, 81% of Rubisco containing Chlamydomonas SSU was located in the condensate. Thus, condensation of Rubisco is strongly associated with Rubisco complexes bearing the Chlamydomonas SSU, which constituted approximately 50% of the Rubisco pool. The latter is consistent with the expected expression levels of plant-algal hybrid Rubisco in S2_Cr_^21^. It is currently unclear if Rubisco can form a heterogenous L8S8 complex with different SSU isoforms, or if only a single SSU isoform is favoured during assembly^26^. Thus, it remains unclear whether the Rubisco pool within the condensate was comprised of a mixture of homogeneous Rubisco complexes, or those containing both Arabidopsis and Chlamydomonas SSUs.

**Figure 3.**
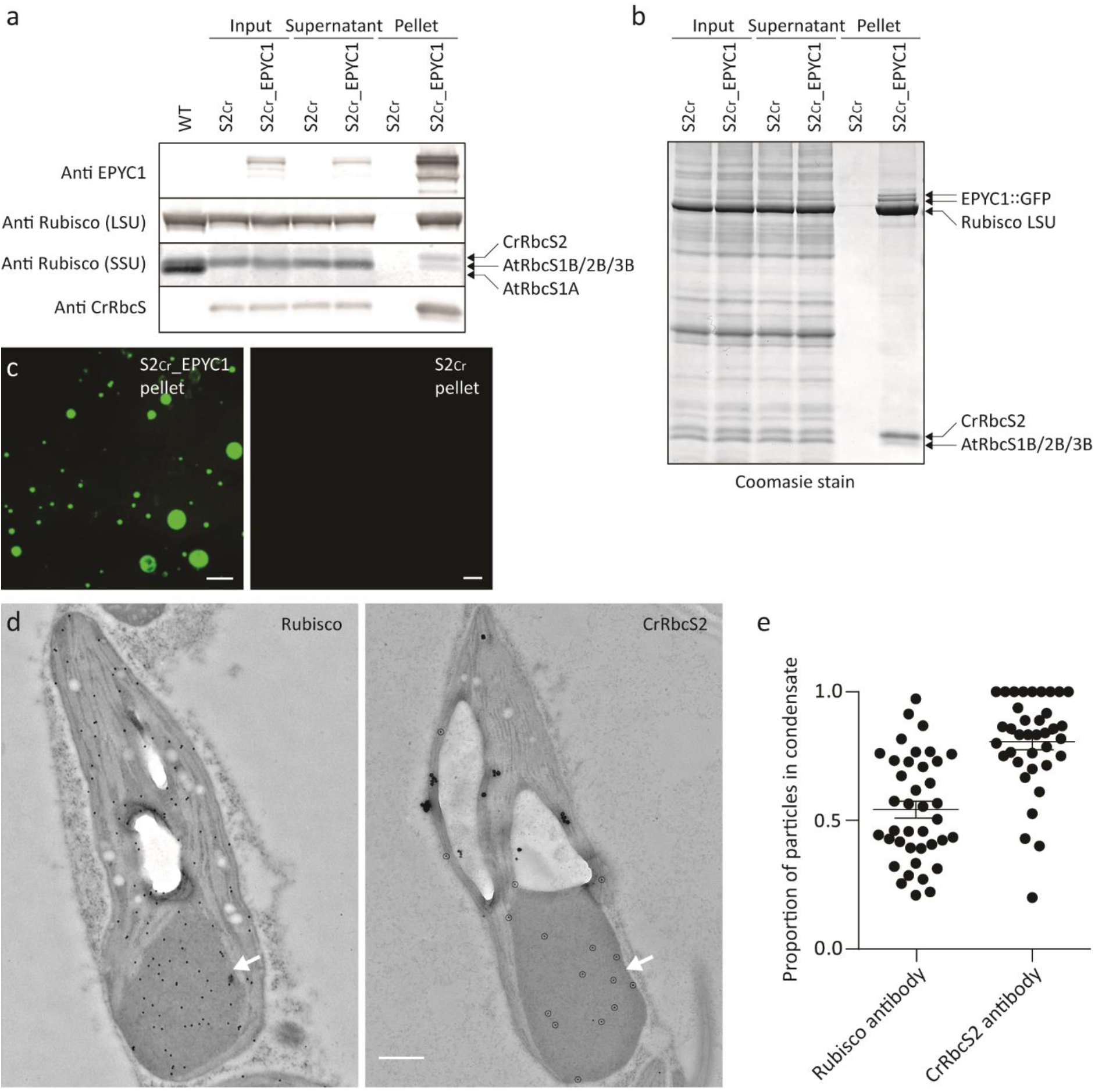
Condensates contain EPYC1 and plant-algal hybrid Rubisco. **a**, EPYC1 and Rubisco protein levels in whole leaf tissue (input), the supernatant following condensate extraction and centrifugation (supernatant) and the pellet (pellet) as assessed by immunoblot analyses with anti-EPYC1, anti-Rubisco (LSU and SSU shown) or anti-CrRbcS2 antibodies. Samples are shown for WT plants (WT), and S2_Cr_ mutants not expressing EPYC1 (S2_Cr_) and expressing EPYC1-dGFP (S2_Cr__EPYC1). The pellet is 40x more concentrated than the input and supernatant. Molecular weights: LSU, 55 kD; RbcS1B, RbcS2B and RbcS3B, 14.8 kD; AtRbcS1A, 14.7kD; CrRbcS2 15.5 kD. **b**, Coomassie-stained SDS-PAGE gel showing the composition of the pelleted condensate. **c**, The condensates in the S2_Cr__EPYC1 pellet coalescence to form large liquid droplets. Scale bar = 50 μm. **d**, Representative immunogold labelling of Rubisco in chloroplasts of an S2_Cr_ transgenic line Ep3 expressing EPYC1-dGFP probed with polyclonal anti-Rubisco (left) or anti-CrRbcS2 (right) antibodies (dots are highlighted for the latter). The condensates are marked by a white arrowhead. Large white structures are starch granules. Scale bar = 0.5 μm. **e**, Proportion of gold nanoparticles inside the condensate compared to the chloroplast for each antibody. The mean and SEM are shown for 37-39 chloroplasts. Scale bar = 0.5 μm.

We compared the growth performance of three separate T2 EPYC1-dGFP S2_Cr_ transgenic lines (Ep1-3) screened for the presence of condensates with their matching T2 azygous segregant S2_Cr_ lines where the EPYC1-dGFP transgene was absent (Az1-3). Plants were cultivated under light levels typical for Arabidopsis growth (200 μmol photons m^-2^ s^-1^) or under higher light levels (900 μmol photons m^-2^ s^-1^) where growth rates are more limited by Rubisco activity (Fig. 4a, 4b, Extended Data Fig. 5)^27^. Regardless of the growth conditions, rosette expansion rates or biomass accumulation were not distinguishable between transformants and their segregant controls. Similarly, T2 EPYC1-dGFP WT plants (EpWT) showed no significant differences compared to T2 segregant lines (AzWT). Due to the reduced Rubisco content in the S2_Cr_ background and differences in Rubisco catalytic characteristics, the growth performance of S2_Cr_ lines was slightly decreased compared to WT plants. The observed differences in growth were in line with those reported previously for S2_Cr_ and WT plants in the absence of EPYC1^21^.

**Figure 4.**
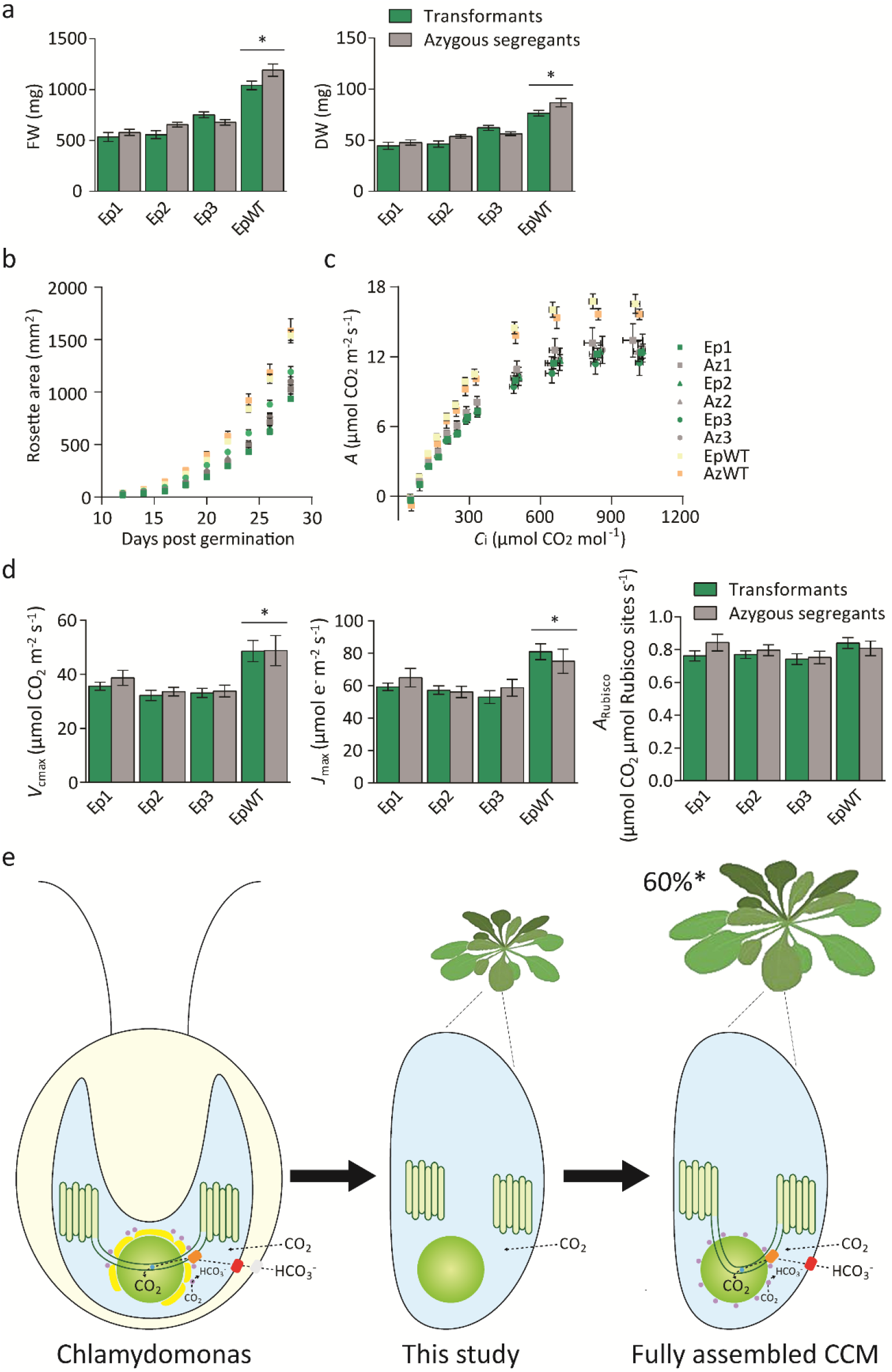
EPYC1-mediated condensation of Rubisco has no negative impact on growth and photosynthesis. **a**, Fresh and dry weights of three T2 EPYC1-dGFP S2_Cr_ transgenic lines (Ep1-3) and an EPYC1-dGFP WT transformant (EpWT) (both in green) with their respective azygous segregants (Az1-3 and AzWT) (in grey). Plants were measured after 32 days of growth under 200 μmol photons m^-2^ s^-1^ light. The mean and SEM are shown for ≥12 individual plants for each line. **b**, Rosette expansion of S2_Cr_ and WT lines in (a). **c**, Net CO_2_ assimilation (*A*)based on intercellular [CO_2_] (*C*_i_) under saturating light (1500 μmol photons m^-2^ s^-1^). Values show the mean and SEM of measurements made on individual leaves from ten or more individual rosettes. **d**, Variables derived from gas exchange data include maximum rate of Rubisco carboxylation (*V*_cmax_), maximum electron transport rate (*J*_max_), stomatal conductance (*G*_s_), mesophyll conductance (*G*_m_) and the net CO_2_ assimilation rate at ambient concentrations of CO_2_ normalized to Rubisco (*A*_Rubisco_). Asterisks indicate significant difference (*P*<0.05) of EpWT lines compared to Ep lines as determined by ANOVA. **e**, Algal CCM components required for enhancing photosynthesis. Generating a pyrenoid-like condensate in a plant chloroplast provides a platform for introducing bicarbonate (HCO_3_^-^) channels/pumps at the chloroplast envelope (e.g. LCIA, shown in red)^33^ and thylakoid membrane (e.g. BST1-3, shown in orange)^35^, a lumenal carbonic anhydrase to convert HCO_3_^-^ to CO_2_ for release into the surrounding Rubisco condensate (CAH3, shown in blue)^35^, mechanisms to capture CO_2_ as HCO_3_^-^ (LCIB and LCIC, shown in purple)^36,37^ and traversing thylakoid membranes^32^. Current models suggest that introduction of a functional biophysical CCM into a C3 plant could lead to productivity gains of up to 60%^8,9,29^.

Photosynthetic parameters derived from response curves of CO_2_ assimilation rate to the intercellular CO_2_ concentration under saturating light were similar between respective Ep and Az lines (Fig. 4c, 4d, Extended Data Table 1). The presence of condensates did not influence the maximum achievable rates of Rubisco carboxylation (*V*_cmax_). Notably, the CO_2_ assimilation rates at ambient concentrations of CO_2_ for Ep and Az lines were comparable to WT lines when normalized for Rubisco content (*A*_Rubisco_) (Fig. 4d). This suggests that the modest reductions in Rubisco turnover rate (*k*_cat_^c^) and specificity (*S*_C/O_) for the plant-algal hybrid Rubisco in S2_Cr_ compared to WT plants have only a mild impact on the efficiency of photosynthetic CO_2_ assimilation and that the observed differences in growth rates are more associated with the reduced levels of Rubisco in S2_Cr_ plants^21^. Mesophyll conductance (*g*_m_) levels were also reduced in all S2_Cr_ lines compared to WT plants, which is consistent with the impact of reduced Rubisco content on *g*_m_ observed in transplastomic tobacco^28^. Condensate formation in the 1A_At_MOD background, where catalytic characteristics of the hybrid Rubisco are indistinguishable from that of WT Rubisco, indicates that the SSU can be further engineered to optimise phase separation, and Rubisco content and performance (Fig. 1c)^19^. Measurements of the maximum electron transport rate (*J*_max_) and the maximum potential quantum efficiency of Photosystem II (*F*_v_/*F*_m_) were also indistinguishable between transformant and segregant lines (Extended Data Table 1). Thus, the apparent displacement of the thylakoid membrane matrix by the condensates (Fig. 1e) had no obvious impact on the efficiency of the light reactions of photosynthesis.

Our findings show that EPYC1 and specific residues on the SSU are sufficient to aggregate Rubisco into a single proto-pyrenoid condensate, which has no apparent negative impact on plant growth. The overall photosynthetic performances of S2_Cr_ transgenic lines appeared unaffected by the condensate, which suggests that conditions inside higher plant chloroplasts are highly compatible with the presence of pyrenoid-type bodies. Our data is arguably the key step for the assembly of a pyrenoid-based CCM into plants that could increase crop yield potentials by >60% (Fig. 4e)^8,9,29^. Previously described approaches for engineering the cyanobacterial carboxysome-based CCM require engineering of the chloroplast-encoded Rubisco large subunit, which is not generally feasible in major grain crops such as wheat and rice^30^, although expression of the large subunit from the nucleus of maize plants has been recently demonstrated^31^. Furthermore, transgenic plants expressing cyanobacterial Rubisco require high concentrations of CO_2_ to grow. Here we demonstrate that condensation of Rubisco is achievable through modification of the nuclear-encoded SSU, which is significantly more amenable to genetic modification. Future work will focus on progressing towards a minimal functional algal CCM by tethering the condensate to a thylakoid membrane for bicarbonate delivery using appropriate transporters^32–34^, as well as introduction of a lumenal carbonic anhydrase and a stromal CO_2_ salvage system^35–37^.

## Methods

### Plant material and growth conditions

Arabidopsis (*Arabidopsis thaliana*, Col-0 background) seeds were sown on compost, stratified for 3 d at 4°C and grown at 20°C, ambient CO_2_ and 70% relative humidity under either 200 or 900 μmol photons m^-2^ s^-1^ supplied by cool white LED lights (Percival SE-41AR3cLED, CLF PlantClimatics GmbH, Wertingen, Germany) in 12 h light, 12 h dark. For comparisons of different genotypes, plants were grown from seeds of the same age and storage history, harvested from plants grown in the same environmental conditions.

### Construct design and transformation

The coding sequence of EPYC1 was codon optimised for expression in higher plants as in Atkinson et al.^19^. Mature EPYC1 was cloned directly into the level 0 acceptor vector pAGM1299 of the Plant MoClo system^38^. To generate fusion proteins, gene expression constructs were assembled into binary level 2 acceptor vectors. The 35S cauliflower mosaic virus (CaMV) promoter and CsVMV (cassava vein mosaic virus) promoter were used to drive expression. Level 2 vectors were transformed into *Agrobacterium tumefaciens* (AGL1) for stable insertion in Arabidopsis plants by floral dipping^39^. Homozygous transgenic and azygous lines were identified in the T2 generation using the pFAST-R selection cassette^40^.

### Protein analyses

Soluble protein was extracted from frozen leaf material of 21-d-old plants (sixth and seventh leaf) in protein extraction buffer (50mM HEPES-KOH pH 7.5 with 17.4% glycerol, 2% Triton X-100 and cOmplete Mini EDTA-free Protease Inhibitor Cocktail (Roche, Basel, Switzerland). Samples were heated at 70 °C for 15 min with 1 x Bolt LDS sample buffer (ThermoFisher Scientific, UK) and 200 mM DTT. Extracts were centrifuged and the supernatants subjected to SDS-PAGE on a 12% (w/v) polyacrylamide gel and transferred to a nitrocellulose membrane. Membranes were probed with rabbit serum raised against wheat Rubisco at 1:10,000 dilution^41^, the SSU RbcS2 from Chlamydomonas (CrRbcS2) (raised to the C-terminal region of the SSU (KSARDWQPANKRSV) by Eurogentec, Southampton, UK) at 1:1,000 dilution, ACTIN (66009-1-Ig, Proteintech, UK) at 1:1000 dilution and EPYC1 at 1:2,000 dilution^6^, followed by IRDye 800CW goat anti-rabbit IgG (LI-COR Biotechnology, Cambridge, UK) at 1:10,000 dilution, and visualised using the Odyssey CLx imaging system (LI-COR Biotechnology).

### qRT-PCR analysis

Total RNA was isolated from leaves of 21-d-old plants (as above) using the RNeasy plant mini kit (Qiagen, USA). Isolated RNA was treated with DNase (Qiagen, USA) and reverse transcribed with random primers using the GoScript Reverse Transcription kit (Promega, USA). Reverse transcription quantitative PCR (RT-qPCR) was carried out with Takyon No ROX SYBR mastermix (Eurogentec, Belgium) as in Atkinson et al.17, and run on a LightCycler 480 (Roche, Switzerland). Gene-specific primers were designed to amplify either eGFP (5’-CAGATTACGCCGTGAGCAAG-3’ and 5’-GCTGAACTTGTGGCCGTTTA-3’) or tGFP (5’-GCATCAGGGGTCTTGAAAGC-3’ and 5’-TCCTTCAAAACGGTGGACCT-3’) (IDT, Belgium). Amplification efficiency was determined with a calibration curve for each primer pair. Two reference genes At4g26410 (RHIP1) and At1g13320 (PP2A) were used for normalization^42^. The relative expression of eGFP and tGFP was calculated according to Vandesompele et al.^43^.

### Condensate extraction

Soluble protein was extracted as before, then filtered through Miracloth (Merck Millipore, Burlington, Massachusetts, USA), and centrifuged at 500 *g* for 3 min at 4 °C, as in Mackinder et al.^6^. The pellet was discarded, and the extract centrifuged again for 12 min. The resulting pellet was washed once in protein extraction buffer, then re-suspended in a small volume of buffer and centrifuged again for 5 min. Finally, the pellet was re-suspended in 25 μl of extraction buffer and used in confocal analysis or SDS-PAGE electrophoresis.

### Growth analysis and photosynthetic measurements

Rosette growth rates were quantified using an in-house imaging system^44^. Maximum quantum yield of photosystem II (PSII) (*F*_v_/*F*_m_) was measured on 32-d-old plants using a Hansatech Handy PEA continuous excitation chlorophyll fluorimeter (Hansatech Instruments Ltd, King’s Lynn, UK)^5^. Gas exchange and chlorophyll fluorescence were determined using a LI-COR LI-6400 (LI-COR, Lincoln, Nebraska, USA) portable infra-red gas analyser with a 6400-40 leaf chamber on either the sixth or seventh leaf of 35-to 45-d-old non-flowering rosettes grown in large pots under 200 μmol photons m^-2^ s^-1^ to generate leaf area sufficient for gas exchange measurements^46^. The response of *A* to the intercellular CO_2_ concentration (*C*_i_) was measured at various CO_2_ concentrations (50, 100, 150, 200, 250, 300, 350, 400, 600, 800, 1000 and 1200 μmol mol^-1^) under saturating light (1,500 μmol photons m^-2^ s^-1^). For all gas exchange experiments, the flow rate was kept at 200 μmol mol^-1^, leaf temperature was controlled at 25 °C and chamber relative humidity was maintained at *ca*. 70%. Measurements were performed after net assimilation and stomatal conductance had reached steady state. Gas exchange data were corrected for CO_2_ diffusion from the measuring chamber as in Bellasio et al.^47^. To estimate *V*_cmax_, *J*_max_, A_Rubisco_, the CO_2_ compensation point (Γ) and mesophyll conductance to CO_2_ (*g*_m_) the *A*/*C*i data were fitted to the C_3_ photosynthesis model as in Ethier and Livingston^48^ using the catalytic parameters *K*_c_^air^ and affinity for O_2_ (*K*_o_) values for wild-type Arabidopsis Rubisco at 25 °C and the Rubisco content of WT and S2_Cr_ lines^21^.

### Confocal laser scanning and fluorescence recovery after photobleaching (FRAP)

Leaves were imaged with a Leica TCS SP8 laser scanning confocal microscope (Leica Microsystems, Milton Keynes, UK) as in Atkinson et al.^17^. Processing of images was done with Leica LAS AF Lite software. Condensate and chloroplast dimensions were measured from confocal images using Fiji (ImageJ, v1.52n)^49^. Condensate volume was calculated as a sphere, while chloroplast volume was calculated as an ellipsoid, where depth was estimated as 25% of the measured width. Chloroplast volumes varied between 24-102 μm^3^, which was within the expected size range and distribution for Arabidopsis chloroplasts^50^. Comparative measurements of pyrenoid area were performed using Fiji on TEM cross-section images of WT *C. reinhardtii* cells (cMJ030) as described in Itakura et al.^23^. FRAP was carried out on live leaf tissue and tissue that had been fixed by infiltrating with 4% (v/v) formaldehyde for 90 min in a vacuum chamber, then infiltrating three times for 5 min each with phosphate-buffered saline (PBS) at pH 7.4. The SP8 microscope was used with a 40x water immersion objective and a photomultiplier tube (PMT) detector. The 488 nm laser was set to 2% power for pre- and post-bleach images, and 25% for the bleaching step. Pre-bleach images were captured at 220 ms intervals (6 in total) and post-bleach images were captured at 400 ms intervals (at least 120). For photo-bleaching the laser was directed to a region with a diameter of 0.5-0.6 μm on one side of the EPYC1 aggregate. Recovery time was calculated by comparing GFP expression to an unbleached region of the same size.

### Super-resolution image microscopy

Super-resolution images were acquired using structured illumination microscopy. Samples were prepared on high precision cover-glass (Zeiss, Jena, Germany). 3D SIM images were acquired on a N-SIM (Nikon Instruments, UK) using a 100x 1.49NA lens and refractive index matched immersion oil (Nikon Instruments). Samples were imaged using a Nikon Plan Apo TIRF objective (NA 1.49, oil immersion) and an Andor DU-897X-5254 camera using a 488nm laser line. Z-step size for z stacks was set to 0.120 μm as required by manufacturer’s software. For each focal plane, 15 images (5 phases, 3 angles) were captured with the NIS-Elements software. SIM image processing, reconstruction and analysis were carried out using the N-SIM module of the NIS-Element Advanced Research software. Images were checked for artefacts using the SIMcheck software (www.micron.ox.ac.uk/software/SIMCheck.php). Images were reconstructed using NiS Elements software v4.6 (Nikon Instruments) from a z stack comprising of no less than 1 μm of optical sections. In all SIM image reconstructions the Wiener and Apodization filter parameters were kept constant.

### Immunogold labelling and electron microscopy

Leaf samples were taken from 21-d-old S2_Cr_ plants and S2_Cr_ transgenic lines expressing EPYC1-dGFP and fixed with with 4% (v/v) paraformaldehyde, 0.5% (v/v) glutaraldehyde and 0.05 M sodium cacodylate (pH 7.2). Leaf strips (1 mm wide) were vacuum infiltrated with fixative three times for 15 min, then rotated overnight at 4°C. Samples were rinsed three times with PBS (pH 7.4) then dehydrated sequentially by vacuum infiltrating with 50%, 70%, 80% and 90% ethanol (v/v) for 1 hr each, then three times with 100%. Samples were infiltrated with increasing concentrations of LR White Resin (30%, 50%, 70% [w/v]) mixed with ethanol for 1 hr each, then 100% resin three times. The resin was polymerised in capsules at 50°C overnight. Sections (1 μm thick) were cut on a Leica Ultracut ultramicrotome, stained with Toluidine Blue, and viewed in a light microscope to select suitable areas for investigation. Ultrathin sections (60 nm thick) were cut from selected areas and mounted onto plastic-coated copper grids. Grids were blocked with 1% (w/v) BSA in TBSTT (Tris-buffered saline with 0.05% [v/v] Triton X-100 and 0.05% [v/v] Tween 20), incubated overnight with anti-Rubisco antibody in TBSTT at 1:250 dilution or anti-CrRbcS2 antibody at 1:50 dilution, and washed twice each with TBSTT and water. Incubation with 15 nm gold particle-conjugated goat anti-rabbit secondary antibody (Abcam, Cambridge, UK) in TBSTT was carried out for 1 hr at 1:200 dilution for Rubisco labelling or 1:10 for CrRbcS2 labelling, before washing as before. Grids were stained in 2% (w/v) uranyl acetate then viewed in a JEOL JEM-1400 Plus TEM (JEOL, Peabody, Massachusetts, USA). Images were collected on a GATAN OneView camera (GATAN, Pleasanton, California, USA).

### Statistical analyses

Results were subjected to analysis of variance (ANOVA) to determine the significance of the difference between sample groups. When ANOVA was performed, Tukey’s honestly significant difference (HSD) post-hoc tests were conducted to determine the differences between the individual treatments (IBM SPSS Statistics Ver. 26.0, Chicago, IL, USA).

## Data availability

All relevant data and plant materials are available from the authors upon request. Raw data corresponding to the figures and results described in this manuscript are available online [*Edinburgh DataShare address to be added*]. Additional data reported in this paper are presented as Extended Data.

## Acknowledgements

This work was funded by the UK Research and Innovation Biotechnology and Biological Sciences Research Council (BB/S015531/1) and Leverhulme Trust (RPG-2017-402). The authors thank Moritz Meyer (Princeton University) for providing additional TEM cross-section images of Chlamydomonas and Ann Wheeler of ESRIC (Edinburgh Super-Resolution Imaging Consortium) for expertise and assistance with SIM. TEM was carried out with the support of Stephen Mitchell and the Wellcome Trust Multi User Equipment Grant (WT104915MA).

---

**Extended Data Table 1.** Photosynthetic parameters from gas exchange and fluorescence measurements for S2_Cr_ transgenic lines of Arabidopsis. The mean and SEM are shown for seven 35-to 45-d-old rosettes for gas exchange variables and twelve 32-d-old rosettes for *F*_v_/*F*_m_. *F*_v_/*F*_m_ is shown for attached leaves dark-adapted for 45 min prior to fluorescence measurements. Letters above the SEM indicate significant difference (P <0.05) as determined by ANOVA followed by Tukey’s HSD tests. Values followed by the same letter are not statistically significantly different. Abbreviations: Γ, CO_2_ compensation point (*C*_i_-*A*); *F*_v_/*F*_m_, maximum potential quantum efficiency of photosystem II; *g*_s_, stomatal conductance to water vapour; *g*_m_, mesophyll conductance to CO_2_ (i.e. conductance of CO_2_ across the pathway from intercellular airspace to chloroplast stroma) using the Ethier and Livingston method^48,52^; *J*_max_, maximum electron transport rate; *V*_cmax_, maximum rate of Rubisco carboxylation.

|  | Ep1 | Az1 | Ep2 | Az2 | Ep3 | Az3 | EpWt | AzWt |
| --- | --- | --- | --- | --- | --- | --- | --- | --- |
| $V_{cmax}$ ($\mu\text{mol CO}_2 \text{ m}^{-2} \text{ s}^{-1}$) | 35.6±1.5<sup>a</sup> | 36.4±2.0<sup>a</sup> | 32.2±1.9<sup>a</sup> | 33.6±1.6<sup>a</sup> | 33.1±1.9<sup>a</sup> | 33.8±2.2<sup>a</sup> | 44.9±1.6<sup>b</sup> | 43.3±1.7<sup>b</sup> |
| $J_{max}$ ($\mu\text{mol e}^- \text{ m}^{-2} \text{ s}^{-1}$) | 59.2±2.3<sup>a</sup> | 61.9±6.3<sup>a</sup> | 57.2±2.6<sup>a</sup> | 56.1±3.5<sup>a</sup> | 52.9±4.4<sup>a</sup> | 58.6±5.2<sup>a</sup> | 76.4±2.4<sup>b</sup> | 74.9±7.5<sup>b</sup> |
| $\Gamma$ ($\mu\text{mol CO}_2 \text{ m}^{-2} \text{ s}^{-1}$) | 63±8<sup>a</sup> | 53±5<sup>a</sup> | 52±6<sup>a</sup> | 54±7<sup>a</sup> | 53±7<sup>a</sup> | 56±8<sup>a</sup> | 51±7<sup>a</sup> | 64±12<sup>a</sup> |
| $g_s$ ($\text{mol H}_2\text{O m}^{-2} \text{ s}^{-1}$) | 0.249±0.031<sup>a</sup> | 0.279±0.051<sup>a</sup> | 0.233±0.017<sup>a</sup> | 0.251±0.015<sup>a</sup> | 0.233±0.021<sup>a</sup> | 0.236±0.016<sup>a</sup> | 0.287±0.018<sup>a</sup> | 0.306±0.011<sup>a</sup> |
| $g_m$ ($\text{mol CO}_2 \text{ m}^{-2} \text{ s}^{-1}$) | 0.034±0.001<sup>b</sup> | 0.035±0.003<sup>b</sup> | 0.032±0.002<sup>b</sup> | 0.033±0.002<sup>b</sup> | 0.034±0.003<sup>b</sup> | 0.032±0.002<sup>b</sup> | 0.045±0.002<sup>a</sup> | 0.046±0.003<sup>a</sup> |
| $F_v/F_m$ (ML) | 0.848±0.002<sup>a</sup> | 0.849±0.002<sup>a</sup> | 0.848±0.001<sup>a</sup> | 0.847±0.001<sup>a</sup> | 0.847±0.002<sup>a</sup> | 0.845±0.002<sup>a</sup> | 0.851±0.002<sup>a</sup> | 0.850±0.001<sup>a</sup> |
| $F_v/F_m$ (HL) | 0.852±0.002<sup>a</sup> | 0.845±0.002<sup>a</sup> | 0.850±0.001<sup>a</sup> | 0.855±0.004<sup>a</sup> | 0.846±0.002<sup>a</sup> | 0.849±0.001<sup>a</sup> | 0.850±0.003<sup>a</sup> | 0.852±0.002<sup>a</sup> |

**Extended Data Figure 1.**
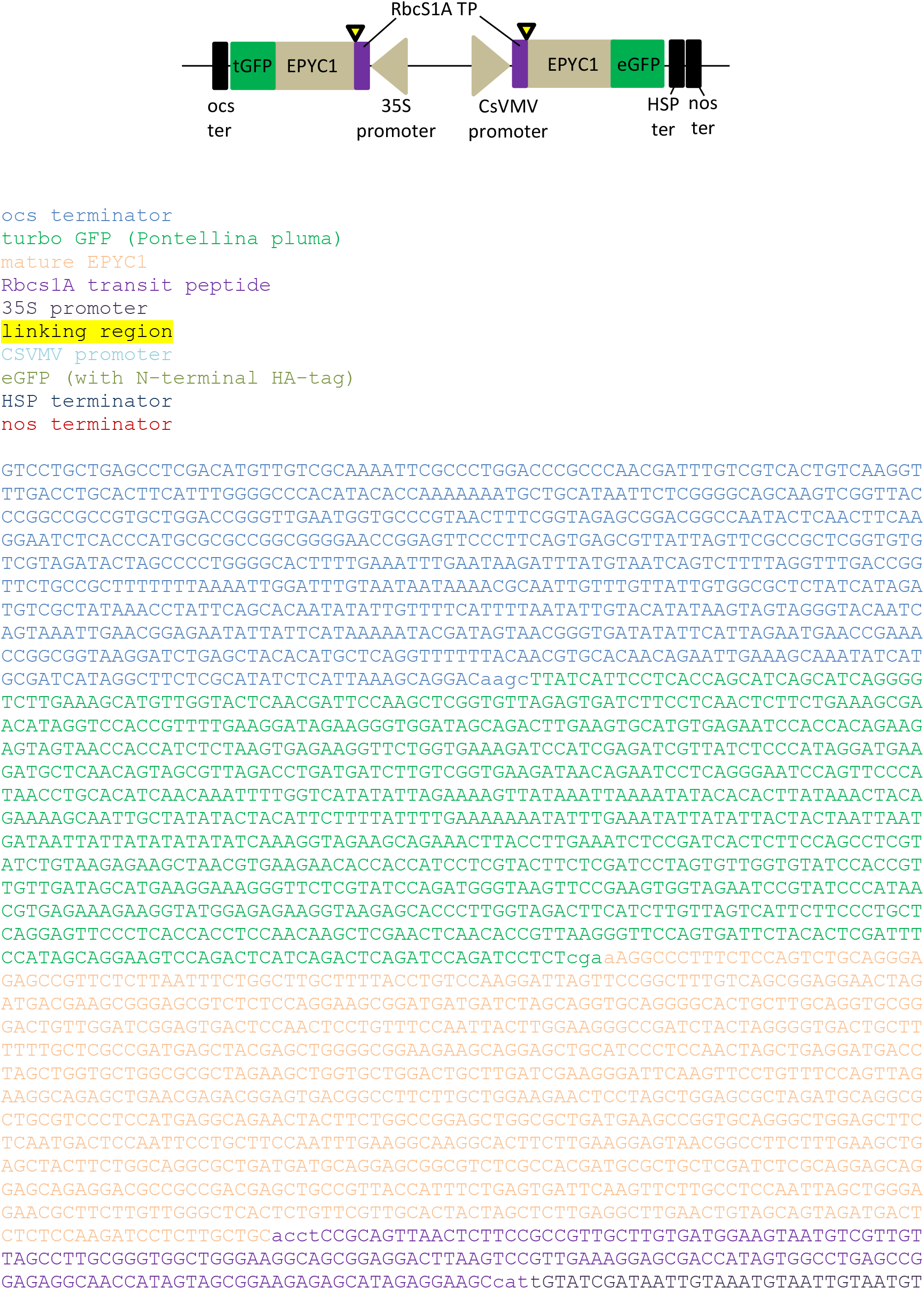
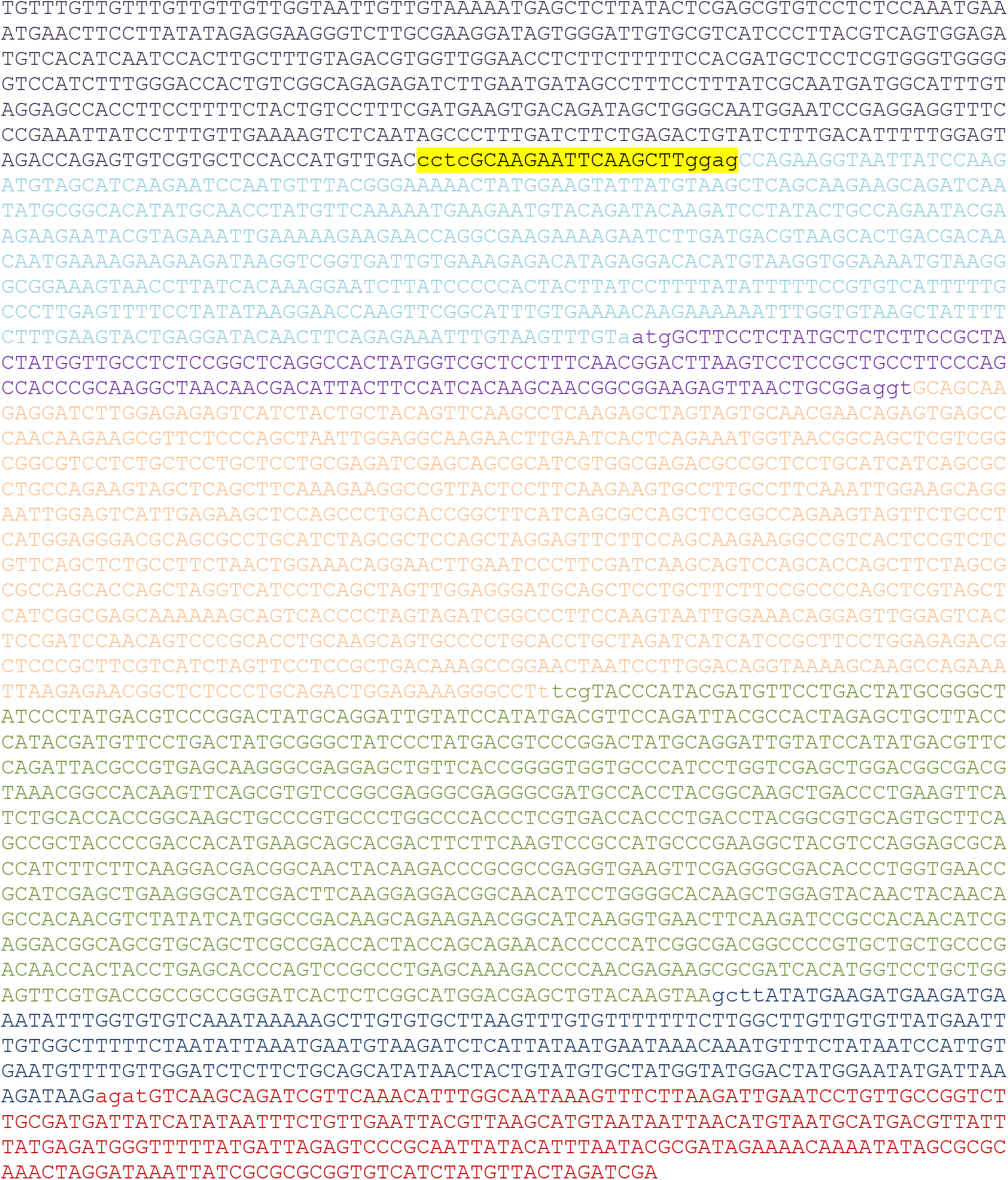
Full sequence of the EPYC1 expression cassettes. Sequence parts are colour coded, and MoClo overhangs are shown in lower case.

**Extended Data Figure 2.**
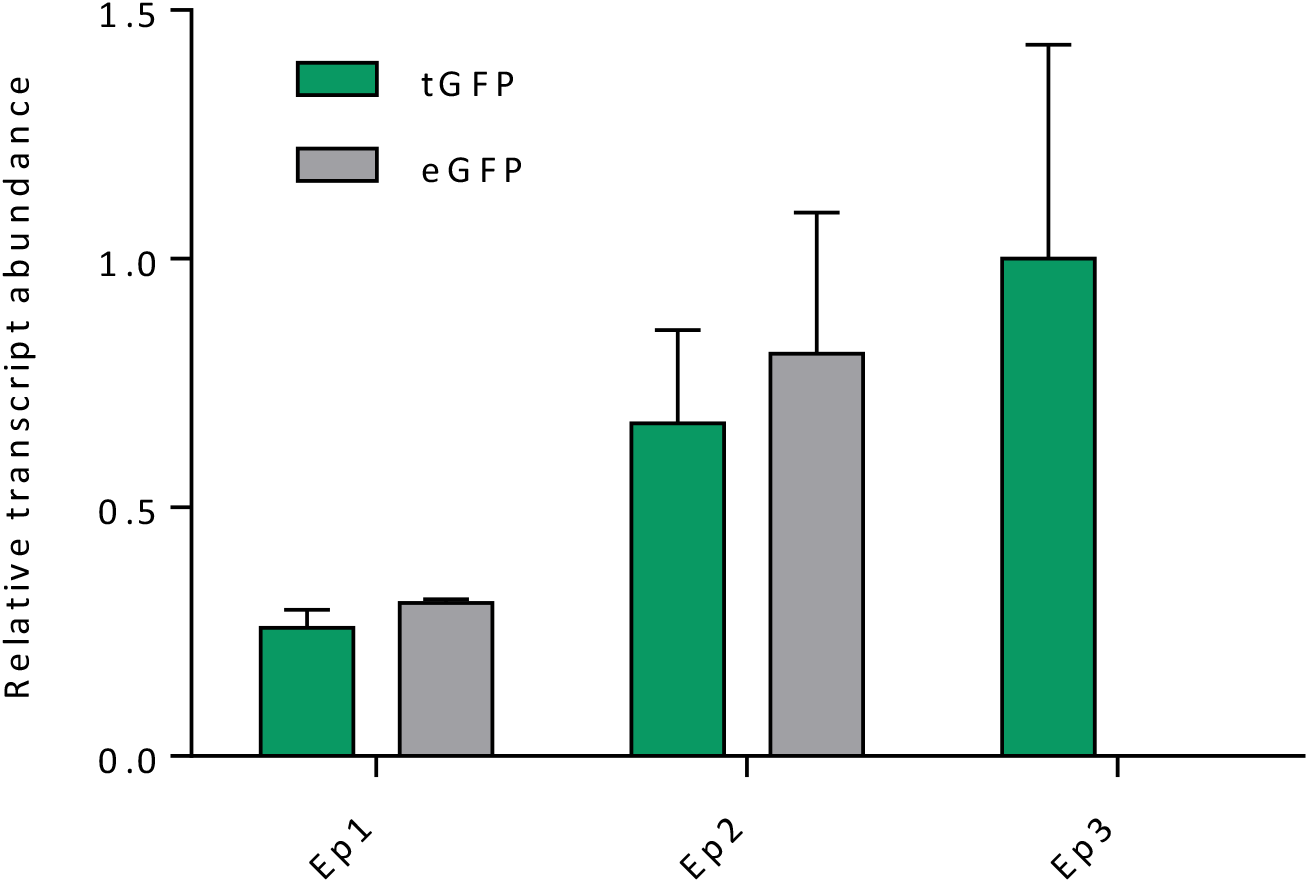
Transcript abundances of tGFP and eGFP in EPYC1-dGFP plant lines. qRT-PCR was carried out on three T2 S2_Cr_ transgenic plants expressing EPYC1-dGFP (Ep1-Ep3, as in Fig. 1B) using gene-specific primers for tGFP and eGFP. Abundances of tGFP and eGFP transcripts are shown relative to the highest tGFP expression level and normalised to reference genes PP2A (At1g13320) and RHIP1 (At4g26410). Error bars show the SEM for two biological replicates.

**Extended Data Figure 3.**
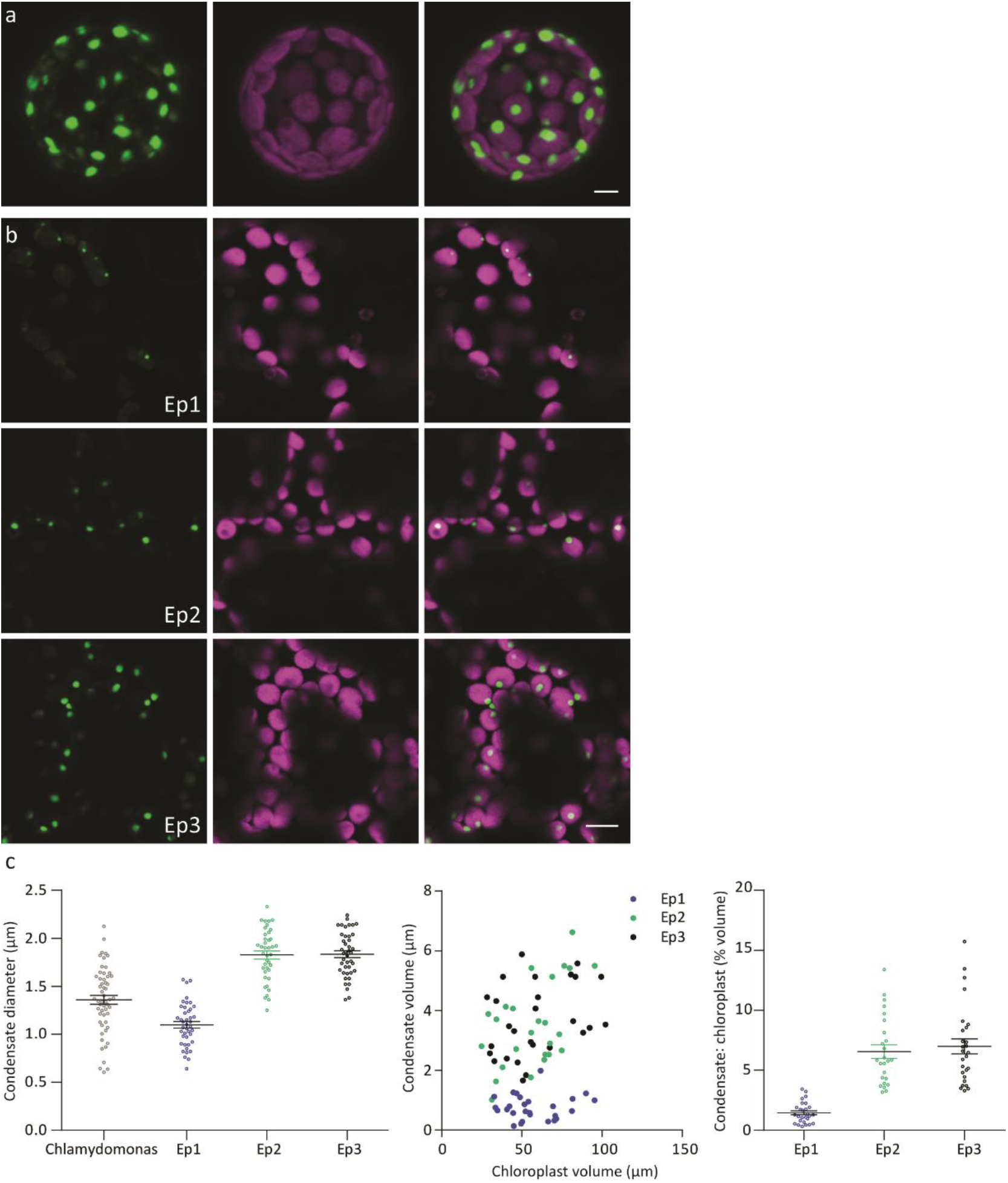
Characterisation of condensates in S2_Cr_ plants. **a**, Maximum projection of a z stack showing condensates in every chloroplast. Scale bar = 5 μm. **b**, The size of condensates is dependent on the expression level of EPYC1. Arabidopsis lines Ep1-3 with different expression levels of EPYC1-dGFP (see Figure. 1b) have different sizes of condensates. Scale bar = 10 μm. **c**, Data derived from confocal images of Chlamydomonas pyrenoids (n=55) and chloroplasts (n=42) from each of the three Ep transgenic lines (Ep1-3) showing the mean diameter and SEM of pyrenoids and condensates (left). The volume of the condensate is shown plotted against estimated chloroplast volume (centre) and the estimated proportion of chloroplast volume occupied by the condensate (right) (n=27 chloroplasts for each line).

**Extended Data Figure 4.**
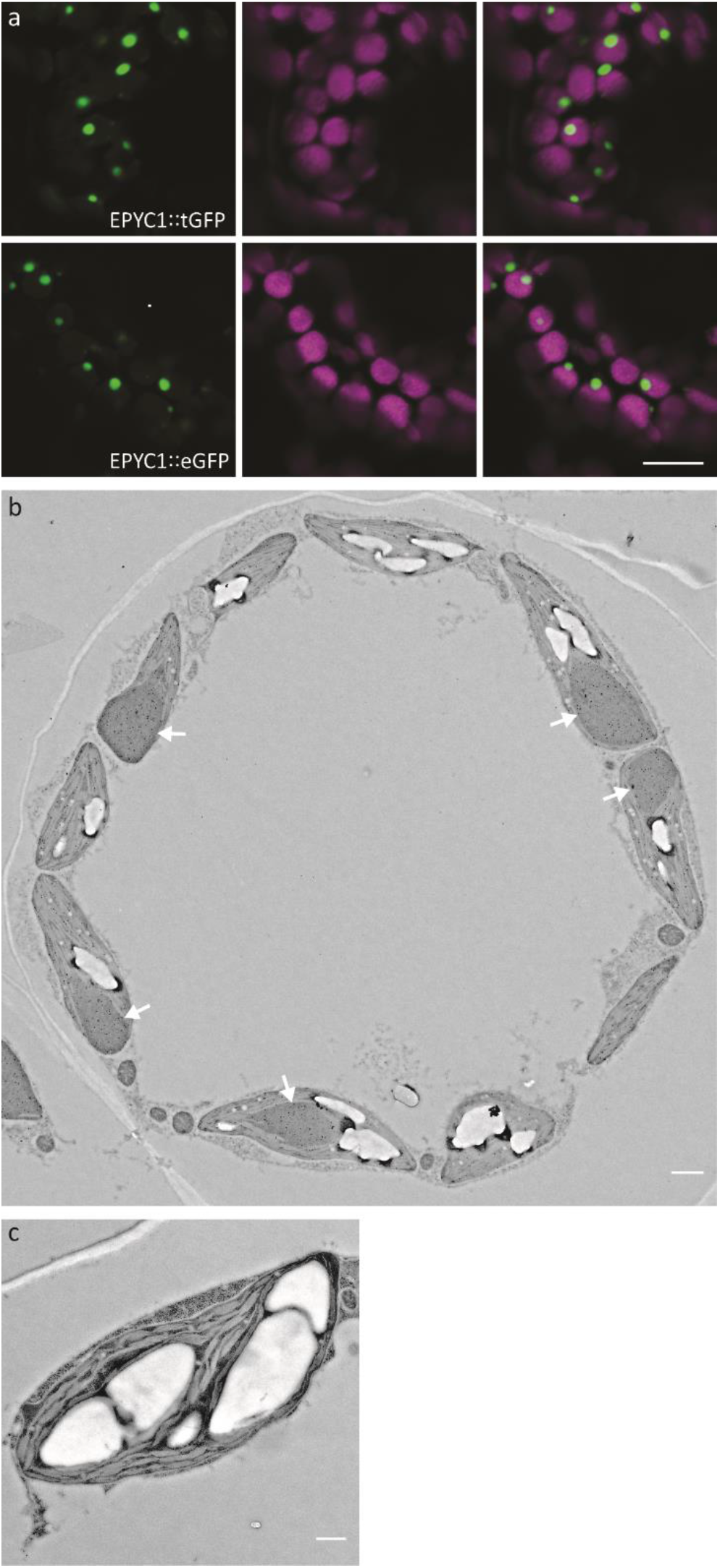
Condensate formation in S2_Cr_ plants. **a**, Condensation occurs in S2_Cr_ plants expressing a single EPYC1 expression cassette. Representative images are shown for EPYC1 fused at the C-terminus to either tGFP (top) or eGFP (bottom). Scale bar = 10 μm. **b**, Representative TEM image of a mesophyll cell cross-section showing chloroplasts with EPYC1-dGFP condensates. Visible condensates are marked by a white arrowhead. The section has been probed by immunogold labelling with anti-Rubisco antibodies. Scale bar = 1 μm. c, Representative TEM image of a chloroplast from a wild-type plant expressing EPYC1-dGFP. No condensates were observed in this background. Scale bar = 0.5 μm.

**Extended Data Figure 5.**
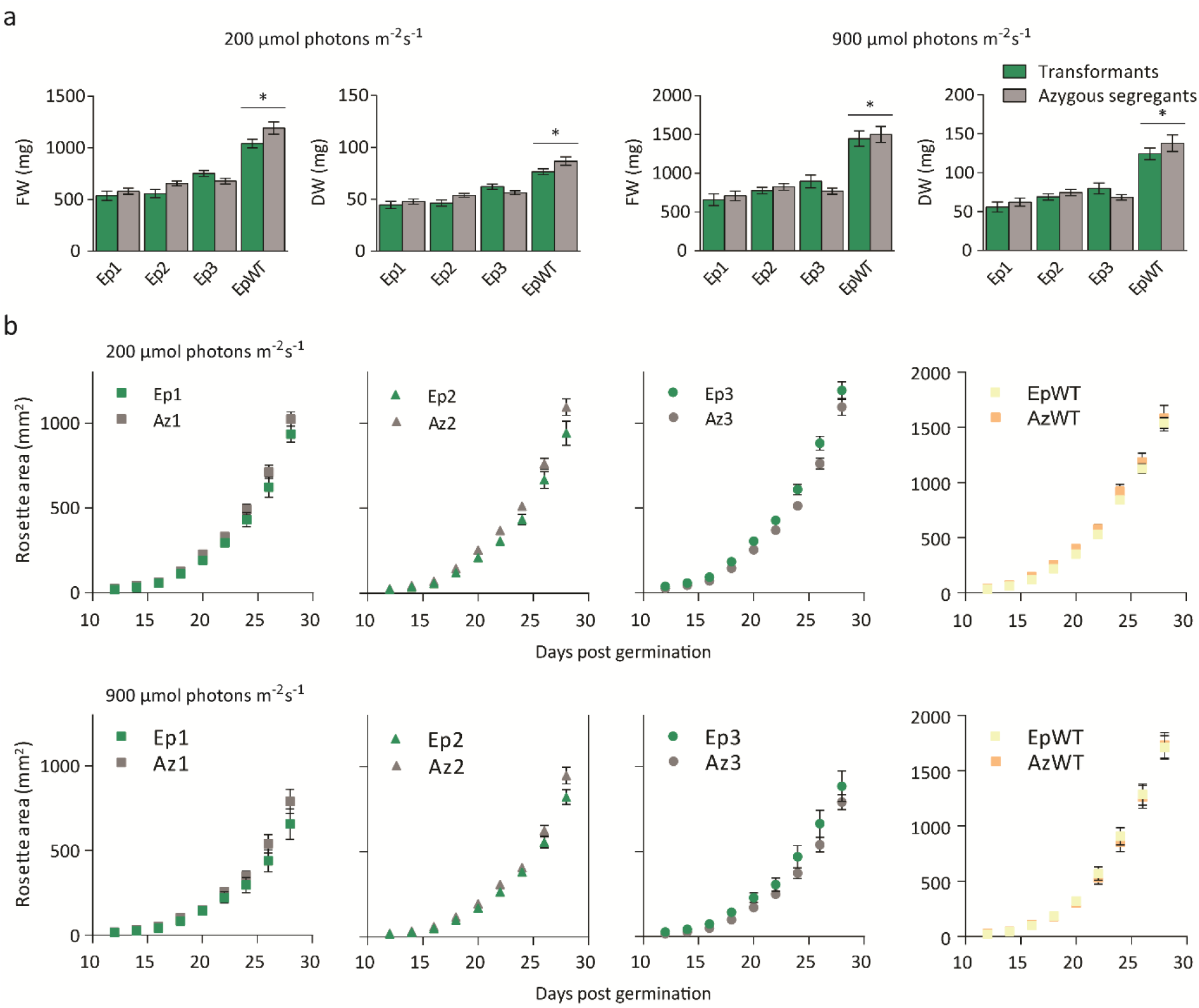
Growth of plant lines expressing EPYC1 under different light levels. **a,** Fresh (FW) and dry weight (DW) of three T2 EPYC1-dGFP S2_Cr_ transgenic lines (Ep1-3) and an EPYC1-dGFP WT transformant (EpWT) with their respective azygous segregants (Az1-3 and AzWT) measured after 32 days of growth at 200 or 900 μmol photons m^-2^ s^-1^ light. Asterisks indicate significant difference (*P*<0.05) as determined by ANOVA. **b,** Rosette area expansion rates for individual transgenic lines and azygous segregants at two different light levels. The mean and SEM are shown for ≥12 individual plants for each line.

